# Deviant cortical sulcation related to schizophrenia, but not cognitive deficits, likely predate brain development in the second trimester

**DOI:** 10.1101/868133

**Authors:** ML. MacKinley, P. Sabesan, L. Palaniyappan

## Abstract

**Objectives:** Aberrant cortical development, inferred from cortical folding measures, is linked to the risk of schizophrenia. Cortical folds develop in a time-locked fashion during fetal growth. We leveraged this temporal specificity of sulcation to investigate the approximate timing of the prenatal insult linked to schizophrenia as well as the cognitive impairment seen in this illness.

**Methods:** Anatomical T1 MRI scans from a publicly available dataset of 68 patients with schizophrenia and 72 controls were used to evaluate the sulcal depth. 5 major primary sulci that are invariable, representing lobar development (calcarine sulcus, superior temporal sulcus, superior frontal sulcus, intraparietal sulcus and inferior frontal sulcus) with formation representing distinct developmental periods were chosen. Sulcal depth was measured using BrainVISA software.

**Results:** A repeated measure Analysis of Variance with 5 sulci and 2 hemispheres as within-subject factors and gender, age and intracranial volume as covariates revealed a significant effect of diagnosis (F[1,134]=14.8, p=0.0002). Control subjects had had deeper superior temporal (left t=3.2, p=0.002; right t=2.8, p=0.006), right inferior frontal (t=2.7, p=0.007) and left calcarine (t=2.2, p=0.03) sulci. A deeper superior frontal sulcus predicted better overall cognitive scores (F[1,54]=8.7, p=0.005) among patients.

**Conclusion:** Our results suggest that the gestational cortical disruption underlying schizophrenia is likely to predate, if not, coincide with the appearance of calcarine sulcus (early 2^nd^ trimester) and affects frontal, temporal and occipital lobes. Nevertheless, the burden of cognitive deficits may relate specifically to aberrant superior frontal development occurring in late 2^nd^ trimester.

## Introduction

An emerging body of evidence implicates aberrations in fetal cortical development to cognitive and mental health outcomes later in life^1,2^. To date, individuals for whom developmental changes in utero are recorded have not been followed up until the emergence of psychosis. Very large prospective cohorts are required for this purpose, given the later emergence and low incident rates of schizophrenia.

The primary sulci of the human brain follow a course of programmed progressive development that occurs in a time-locked fashion and is highly sensitive to foetal disruptions^1^. The development of the primary sulci defining lobar development (calcarine sulcus, superior temporal sulcus, superior frontal sulcus, intraparietal sulcus and inferior frontal sulcus) is so precise in developing foetuses (with emergence at 16, 23, 25, 26 and 28 weeks respectively^1^) that these structures can be used to estimate gestational age and brain maturation^2^. Studying the location of aberrations in cortical folding in adult life can serve as a window to the time-locked disruptions suffered by the developing cortical architecture^3^.

Clinically, atypical development in the primary sulci has been associated with sensory and information processing deficits among patients with schizophrenia^4–8^, potentially contributing to the development of both cognitive deficits as well as psychotic symptoms. The likely timing of embryonic/fetal disruption in schizophrenia continues to be unknown, with prenatal immune models implicating both second and third trimester insults as risk factors ^9^, while maternal stress and malnourishment models implicate third trimester insults ^9,10^. Prenatal insults not only associate with risk of schizophrenia, but also influence the severity of the eventual illness (e.g. the presence of cognitive deficits ^11^). We compared sulcal depth of 5 major primary sulci – calcarine, superior temporal, superior frontal, intraparietal, and inferior frontal- that are anatomically district, with temporal specificity in their gestational appearance based on Chi et al.^1^, among healthy controls and schizophrenia patients and assessed their relationship to cognitive performance in patients.

## Method

68 patients with schizophrenia or schizoaffective disorder and 72 healthy controls from the National Institute of Health Centre for Biomedical Research Excellence (COBRE) grant were included in the analysis. Patients were excluded based on history of neurological disorder, intellectual disability, severe head trauma, or substance abuse or dependence within the last 12 months. Diagnostic information was collected using the Structured Clinical Interview used for DSM Disorders (SCID)^12^. Seven domains of cognition (speed of processing, attention/vigilance, working memory, verbal learning, visual learning, reasoning/problem solving, and social cognition) were assessed using the Measurement and Treatment Research to Improve Cognition in Schizophrenia (MATRICS) battery^18^.

MRI data was collected on a Siemens 3T TIM Trio scanner. A multi-echo MPRAGE (MEMPR) sequence was used^13^. Sulcal depth was measured using Morphologist interface of BrainVISA 4.5 with default settings. Following the construction of 3-dimensional models of cortical folds, various sulci were automatically classified using a probabilistic algorithm with maximum depth computed for each identified sulcus. The identified sulci were visually inspected by 2 raters to ensure that the boundaries are in accordance with Ono’s Atlas of Cerebral Sulci (Ono et al., 1990). Repeated measures ANOVA were used with the 5 sulci of interest and two hemispheres as within-subject factors and gender, age, and intracranial volume as covariates to assess for differences between HCs and patients. 5 non-collinear factors representing the 5 bilateral sulci were obtained using varimax rotation and related to overall MATRICS standardized composite score in patients using multiple regression.

## Results

No statistically significant differences on gender, age, handedness or intracranial volume between patients and controls were identified, but as expected patients had lower scores on the MATRICS composite score, as well as several specific cognitive domains including processing speed, attention vigilance, working memory, and problem solving (Table 1).

**Table 1:** Group differences in cognitive and demographic features.

| **Variable** | **Healthy Control**<br>**n=72** | **Patients**<br>**n=68** |
| --- | --- | --- |
| **Gender (M/F)** | 51/21 | 55/13 |
| **Age [ m (sd) ]** | 35.87 (11.74) | 38.49 (13.86) |
| **Handedness (Right / Left / Both)** | 69/1/2 | 57/9/2 |
| **Intracranial Volume in mL [ m (sd)]** | 1511.26 (141.58) | 1543.51 (157.97) |
| **MATRICS Composite Score** | 49.97 (8.31) | 32.02 (13.42)* |
| Processing Speed [ m (sd) ] | 54.12 (8.94) | 35.48 (12.21) * |
| Attention Vigilance [ m (sd) ] | 48.57 (9.23) | 37.56 (13.37) * |
| Working Memory [ m (sd) ] | 50.86 (10.22) | 40.84 (12.84)* |
| Verbal Learning [ m (sd) ] | 45.88 (8.98) | 38.91 (8.89) |
| Visual Learning [ m (sd) ] | 45.25 (10.16) | 35.96 (11.97) |
| Problem Solving [ m (sd) ] | 55.37 (8.44) | 44.50 (11.55) * |
| Social Cognition [ m (sd) ] | 51.46 (10.04) | 41.91 (10.46) |

The ANOVA showed a significant between-subjects effect for diagnosis (F[1,134]=14.8, p=0.0002), with significant effects also identified for gender (F[1,134]=7.4, p=0.007, sulcal depth for females>males) and age (F[1,134]=4.5, p=0.035; reduced depth with increasing age). Parameter estimates revealed a significant effect of diagnosis (Controls>Patients) for left superior temporal (t=3.2, p=0.002), right superior temporal (t=2.8, p=0.006), right inferior frontal (t=2.7, p=0.007) and left calcarine (t=2.2, p=0.03) sulci. As age and gender had significant effects on the model, we used them as covariates for the multiple regression analysis relating MATRICS total score to the 5 orthogonal sulcal factor scores. The depth of the superior frontal sulcus was the only predictor of the variation in the cognitive score (t=2.88, p=0.006; all other predictors p>0.17; overall model F[7,52]=2.15, p=0.055). Patients with a deeper superior frontal sulcus had higher composite cognitive scores in patients. A similar predictive model for MATRICS composite scores in healthy controls was not significant (overall model F[7,49]=1.56, p=0.17).

## Discussion

These findings suggest that developmental disruptions predominantly affect morphology of the frontal, temporal and occipital lobes cortical disruption in patients with schizophrenia, with onset likely to predate or coincide with the appearance of calcarine sulcus (i.e. 16 weeks, early 2^nd^ trimester). In contrast to illness-related developmental deviation, the burden of cognitive deficits seen among patients may relate specifically to aberrant superior frontal development occurring in late 2^nd^ trimester.

**Figure 1:**
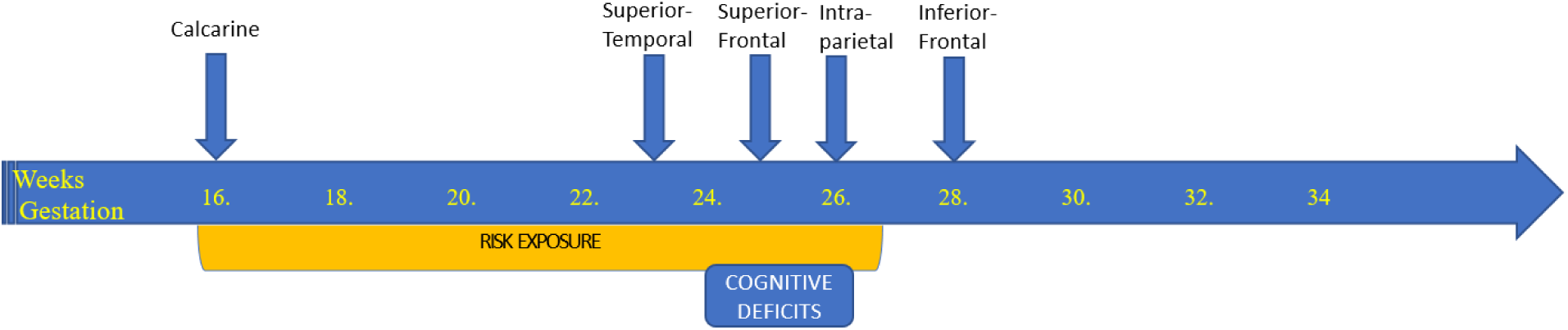
Timeline of the emergence of the 5 primary sulci under study, and the posited risk exposure.

While sulcal depth is fairly constant in early adult life, age-related changes have been reported in adolescence [0.3%/yr. age 11-17] ^14^and later life, especially affecting the superior frontal sulcus^15^. These changes are considerably smaller in magnitude (<10 times) when compared to age-related changes in sulcal width^16^ and other morphometric features. We had age-matched case-control groups and adjusted for age in our regression analysis. Our observations relating schizophrenia to earlier aberrations in development (∼16wks), while the cognitive deficits of schizophrenia to later development (∼25 wks) raises an interesting question whether transient but time-locked insults to the developing brain are pathogenic, while sustained or repeated insults (that extend to later part of development), lead to a more severe form of illness characterised by cognitive deficits. Given the low genetic correlation between cognitive impairment and schizophrenia, our results support the notion that the development of these two features may be somewhat independent ^17^, at least on the timescale of cortical development. Relevant animal models of in utero risk exposure are required to address this question.

## Acknowledgements

Data was downloaded from the Collaborative Informatics and Neuroimaging Suite Data Exchange tool (COINS; http://coins.mrn.org/dx) and data collection was performed at the Mind Research Network, and funded by a Center of Biomedical Research Excellence (COBRE) grant 5P20RR021938/P20GM103472 from the NIH to Dr. Vince Calhoun.

